# Contrasting population structure and demographic history of cereal aphids in different environmental and agricultural landscapes

**DOI:** 10.1101/2020.01.16.909077

**Authors:** Ramiro Morales-Hojas, Jingxuan Sun, Fernando Alvira Iraizoz, Xiaoling Tan, Julian Chen

## Abstract

Genetic diversity of populations has important ecological and evolutionary consequences, which are fundamental to improve the sustainability of agricultural production. Studies of how differences in agricultural management and environment influence the population structure of insect pests are fundamental to predict outbreaks and optimise control programmes. Here, we have studied the population genetic diversity and evolution of *Sitobion avenae* and *Sitobion miscanthi* (previously mistaken for *S. avenae*), which are among the most relevant aphid pests of cereals across Europe and China, respectively. We have used a genomic approach that allows the identification of weak geographic structure and migration patterns at scales that were previously not discernible. In the present study, we show that the population structure in present day populations are different from that described in previous studies, which suggests that they have evolved recently possibly as a response to human-induced changes in agriculture. In the UK, *S. avenae* is predominantly anholocyclic and, as a result of the evolution of insecticide resistance, a superclone is now dominant across the geographic distribution in the country and the genetic diversity is low. In China, *S. miscanthi* populations are mostly holocyclic, with one sexual stage in autumn to produce overwintering eggs, and there are six genetically differentiated subpopulations and high genetic differentiation between geographic locations, which suggests that further taxonomical research is needed. Unlike in the case of *S. avenae* in England, there is no evidence for insecticide resistance and there is no predominance of a single lineage in *S. miscanthi* in China.

## Introduction

A major challenge in agricultural entomology is to develop efficient control strategies for pest organisms. For this, it is important to understand how environmental and anthropogenic factors influence the genetic structure and the evolutionary dynamics of insect populations, both for beneficial and pest species. The level of genetic structure and diversity is the result of a combination of several factors which include selection, migration and life history (i.e. reproduction mode), and studying their consequences on insect populations is of great interest to improve ecological agricultural practices. The use of pesticides remains a necessary way to control and manage pests in agriculture. However, their use imposes a strong selection pressure on pest populations and resistance to different types of insecticides has, therefore, evolved in many insects (Bass, Denholm, Williamson, & Nauen, 2015; Georghiou, 1972). Designing new strategies of pest control that rationalise the use of insecticides and reduce the likelihood of an evolution of resistance has become key for the development of sustainable agriculture practices that reduce the environmental footprint. Understanding how pest populations respond to selective pressure and adapt to ecological changes is key to design rational strategies of management and control that are more targeted. In addition, a better understanding of the geographic connectivity between populations and the dispersal capacity of pests provides valuable information to control their abundance and distribution, while preventing also the spread of adaptive genetic variation, such as insecticide resistance, across their geographic range. Therefore, it is essential that we incorporate the fundamental knowledge about population genetics into agricultural entomology.

Aphids comprise some of the most pernicious species of crop pests. In cereals, *Sitobion avenae* and *Sitobion miscanthi* are two of the most economically important species in Europe and Asia, respectively, and they are major vectors of the barley yellow dwarf virus (BYDV), which can severely reduce cereal yield (Vickerman & Wratten, 1979). Both *Sitobion* species are monoecious, feeding only on Poaceae grasses and cereals. Like many aphids, *S. avenae* and *S. miscanthi* show different levels of variation in the life-cycle, from individuals that are obligate cyclical parthenogenetic and have a generation that undergoes sexual reproduction in the primary host (holocycly), to clones that are obligate parthenogenetic and reproduce asexually all year round (anholocycly) (Dedryver, Le Gallic, Gauthier, & Simon, 1998). In addition, individuals can remain asexual in the cereal crops during winter as a response to environmental cues such as warmer temperatures and day length. As a result, a geographic cline in the reproductive type has been described in the *S. avenae* populations of UK and France, with increasing proportion of sexual reproduction towards the north of the countries (Llewellyn et al., 2003; Simon et al., 1999). In the case of *S. miscanthi*, variation in the life-cycle has also been described. Populations from this species in Australia and New Zealand are anholocyclic, while they are holocyclic in Taiwan (Sunnucks, England, Taylor, & Hales, 1996; Wilson, Sunnucks, & Hales, 1999). In China, *S. miscanthi* has been traditionally reported to be anholocyclic (Guo, Shen, Li, & Gao, 2005; Zhang, 1999). However, contrary to the observations in *S. avenae* and other species, a recent population genetics study has observed signatures of cyclical parthenogenesis in the southern populations of the country while obligate parthenogenetic reproduction would be dominant in the north (Wang, Hereward, & Zhang, 2016).

Resistance to pyrethroids was first detected in the UK populations of *S. avenae* in 2011. This was due to a knockdown resistance (*kdr*) mutation (L1014F) in the sodium channel gene (Foster et al., 2014). This mutation appeared in heterozygosity in one clone of *S. avenae*, known in the literature as clone SA3, and rapidly increased its abundance in the UK population from 2009 to 2014, although in variable proportions in different locations and years (Dewar & Foster, 2017; Malloch, Foster, & Williamson, 2016; Malloch, Williamson, Foster, & Fenton, 2014). The spread of the mutation in the UK was limited by the fact that the SA3 clone is anholocyclic, so pyrethroid resistance has not spread to other lineages through sexual recombination. In addition, the high connectivity of the UK populations (Llewellyn et al., 2003), probably facilitated the geographic spread of the resistant clone from its location of origin. Therefore, the continued use of pyrethroids combined with the long dispersal capacity of *S. avenae* has likely favoured the spread of this clone across the UK. Nevertheless, the clonal diversity remained high and similar to the diversity before the evolution of pyrethroid resistance, and other susceptible clones and phenotypes were still present in different proportions in British populations by 2015 (Llewellyn et al., 2003; Malloch et al., 2016). In the case of *S. miscanthi*, there is no available information in the literature regarding the evolution of insecticide resistance. However, understanding the dynamics and movement of the species can help manage and control the damage in cereal crops, and establish management programs to reduce the likelihood of insecticide resistance evolution. In China, previous studies have shown high levels of genetic diversity in *S. miscanthi* (Guo et al., 2005; Wang et al., 2016), similar to those reported for *S. avenae* in the UK and France, and there is genetic differentiation between north and south of the country but low differentiation within each region, suggesting free gene flow within geographic regions (Guo et al., 2005; Wang et al., 2016).

In the present study we analyse the population genetics and demographic history of *S. avenae* and *S. miscanthi* in England and China, respectively, using a genomic approach to identify potential differentiation at a fine scale. We discuss the results in view of the differences in life-history types and the evolution of insecticide resistance, which may be limited by the reproductive type. These genomics approaches have identified genetic variation at a national and regional scale for other aphids in regions where they disperse long distances (Morales-Hojas et al., 2019).

## Materials and Methods

### Samples

Individuals of *S. avenae* were collected during June-July 2018 using the 12 English 12.2 m high suction-traps ran by Rothamsted Insect Survey (Table 1). Individuals of *S. miscanthi* were collected in 10 sites across the cereal growing areas of China (Table 1). The 10 individuals of *S. miscanthi* were collected from the same wheat field but from plants separated by 10 m to reduce the probability of sampling the same clone.

**Table 1.**
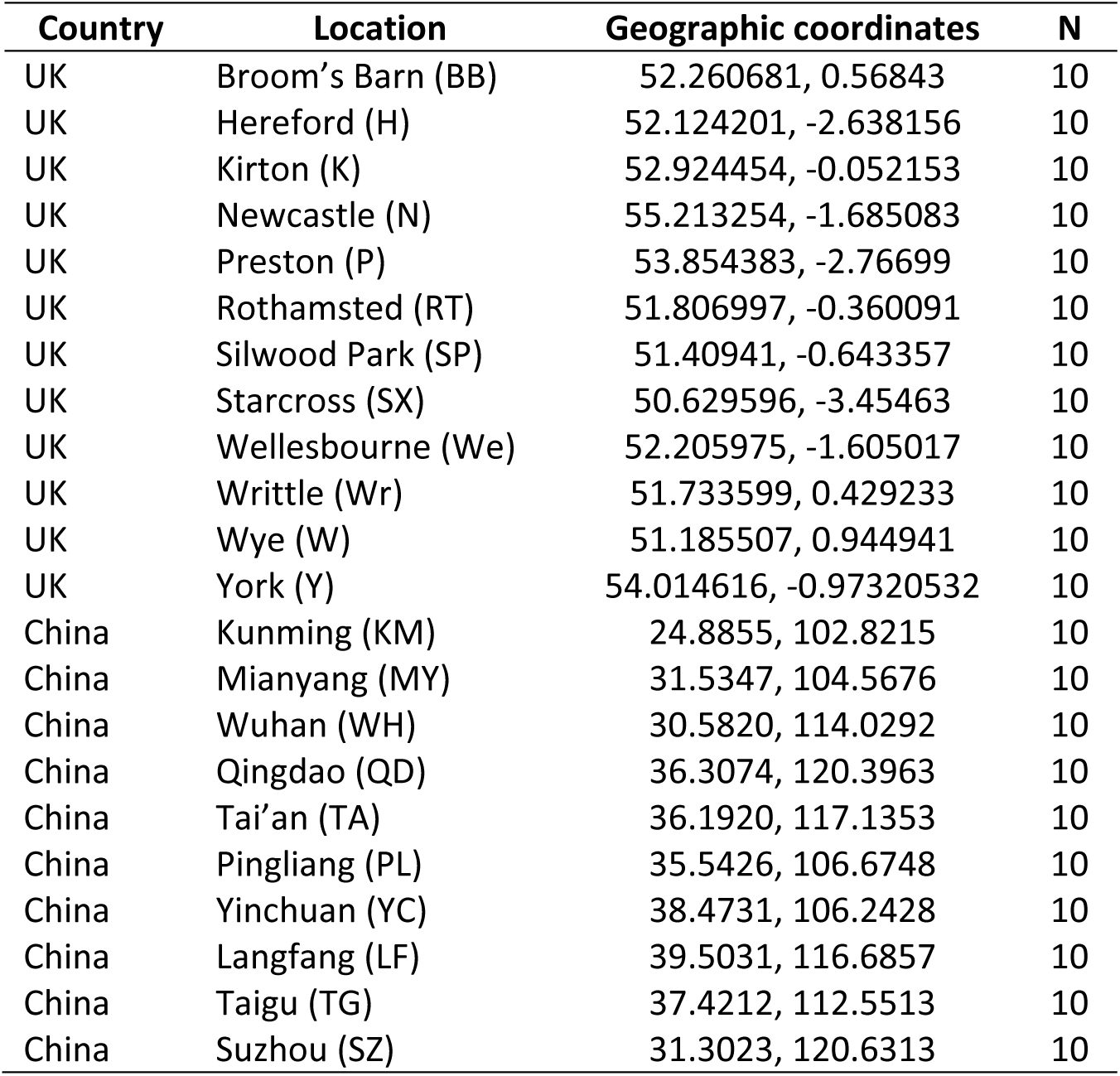
Locations and number of samples (N) used in the present study.

| Country | Location | Geographic coordinates | N |
| --- | --- | --- | --- |
| UK | Broom's Barn (BB) | 52.260681, 0.56843 | 10 |
| UK | Hereford (H) | 52.124201, -2.638156 | 10 |
| UK | Kirton (K) | 52.924454, -0.052153 | 10 |
| UK | Newcastle (N) | 55.213254, -1.685083 | 10 |
| UK | Preston (P) | 53.854383, -2.76699 | 10 |
| UK | Rothamsted (RT) | 51.806997, -0.360091 | 10 |
| UK | Silwood Park (SP) | 51.40941, -0.643357 | 10 |
| UK | Starcross (SX) | 50.629596, -3.45463 | 10 |
| UK | Wellesbourne (We) | 52.205975, -1.605017 | 10 |
| UK | Writtle (Wr) | 51.733599, 0.429233 | 10 |
| UK | Wye (W) | 51.185507, 0.944941 | 10 |
| UK | York (Y) | 54.014616, -0.97320532 | 10 |
| China | Kunming (KM) | 24.8855, 102.8215 | 10 |
| China | Mianyang (MY) | 31.5347, 104.5676 | 10 |
| China | Wuhan (WH) | 30.5820, 114.0292 | 10 |
| China | Qingdao (QD) | 36.3074, 120.3963 | 10 |
| China | Tai'an (TA) | 36.1920, 117.1353 | 10 |
| China | Pingliang (PL) | 35.5426, 106.6748 | 10 |
| China | Yinchuan (YC) | 38.4731, 106.2428 | 10 |
| China | Langfang (LF) | 39.5031, 116.6857 | 10 |
| China | Taigu (TG) | 37.4212, 112.5513 | 10 |
| China | Suzhou (SZ) | 31.3023, 120.6313 | 10 |

### DNA extraction and SNP genotyping

DNA was extracted from samples using Qiagen’s DNeasy Blood and Tissue kit following the manufacturer’s protocol. Samples of *S. avenae* and *S. miscanthi* were genotyped separately. Genomic DNA was digested with *MseI* in the case of *S. avenae* and with *ApeKI* in the case of *S. miscanthi*. The library preparation was performed following the standard Illumina pair-end (PE) protocol, and PE sequencing of 150 bp was performed on an Illumina HiSeq platform. Library preparation and sequencing of samples were outsourced commercially. Reads quality was assessed with FastQC v0.67 and, in the case of *S. miscanthi*, the first 10 bases were trimmed due to low quality using trimmomatic 0.36.1 (Bolger, Lohse, & Usadel, 2014). Reads were mapped to a draft of the *S. avenae* genome. Duplicates were removed using MarkDuplicates v2.7.1.1 and indels were realigned with BamLeftAlign v1.0.2.29-1. Variant calling was carried out with FreeBayes v1.0.2.29-3 (Garrison & Marth, 2012). The resulting SNPs from FreeBayes were annotated using snpEff v4.0. These tools were run using Galaxy v17.05 (Afgan et al., 2016). SNPs called with FreeBayes were filtered using VCFtools v0.1.14 (Danecek et al., 2011) before the markers were used in subsequent analyses. Different filtering schemes were used to obtain a dataset that maximised the quality of the SNPs and genotypes while minimising the missing data at marker and individual levels (Supplementary Table S1), as recommended by O’Leary, Puritz, Willis, Hollenbeck, and Portnoy (2018).

### Analyses of population structure

The population structure of both species was investigated using the Bayesian genetic clustering algorithm implemented in Structure 2.3.4 (Pritchard, Stephens, & Donnelly, 2000). We used the admixture model with correlated frequencies and, to detect any potential subtle genetic structure, we ran Structure with the sampling locations set as priors (locprior = 1); this model has the power to detect a weak structure signal and does not bias the results towards detecting genetic structure when there is none. In the case of *S. miscanthi*, a first run of Bayesian clustering analyses of the population structure was carried out with 5 independent simulations with 100,000 burn-in and 100,000 mcmc chains for each of *K* 1–10. An additional run of 5 independent simulations with 100,000 burn-in and 500,000 mcmc chains was carried out for *K* 5–10 to confirm the results of the first run and ensure convergence in the mcmc step. In the analyses of *S. avenae* from the UK, Structure was run with 5 replicates of 500,000 burn-in and 1,000,000 mcmc chains for *K* ranging from 1 to 12. Parameter convergence was inspected visually. We ran the Structure simulations using a multi-core computer with the R package ParallelStructure (Besnier & Glover, 2013) in the CIPRES science gateway server (Miller, Pfeiffer, & Schwartz, 2010). The number of *K* groups that best fitted the dataset was estimated using the method of Evanno, Regnaut, and Goudet (2005) using Structure Harvester Web v0.6.94 (Earl & Vonholdt, 2012). Cluster assignment probabilities were estimated using the programme Clumpp (Jakobsson & Rosenberg, 2007) as implemented in the webserver CLUMPAK (Kopelman, Mayzel, Jakobsson, Rosenberg, & Mayrose, 2015).

The genetic diversity measures of the populations were estimated using Arlequin 3.5.2.2 (Excoffier, Laval, & Schneider, 2005). Genetic variation among populations was investigated using an Analysis of the Molecular Variance (AMOVA) with 10000 permutations using Arlequin 3.5.2.2. We used hierarchical AMOVA to test the population structures resulting from the Structure runs. Population pairwise divergence was investigated using F_ST_, and the significance was evaluated with 10000 permutations in Arlequin. We ran a Mantel test as performed in Arlequin to evaluate the correlation between the genetic distances (F_ST_) and the geographic distance between sampling locations in China estimated using Google maps (Supplementary Table S2). The demographic history of *S. avenae* was also explored using Arlequin. For this, we first estimated the gametic phase from the multilocus diploid data using the ELB algorithm with the default parameter values. Population expansion or bottleneck were inferred using Fu’s F_S_ (Fu, 1997), and mismatch analyses were run with 1000 bootstrap replicates to estimate the Harpending’s raggedness index and the sum of squared deviation (SSD) given the population expansion and spatial expansion models.

Phylogenetic trees for *S. avenae* were constructed using Maximum Likelihood (ML) with RaxML 8.2.12 (Stamatakis, 2014) run in the server CIPRES (Miller et al., 2010). RAxML was run with 1000 bootstrap inferences with subsequent ML search using the gtrgamma model. The Lewis correction for ascertainment bias was implemented as it is the appropriate model for binary datasets that include only variable sites (as it is the case of SNPs) (Leache, Banbury, Felsenstein, de Oca, & Stamatakis, 2015; Lewis, 2001).

## Results

### *Genetic diversity and population structure of* S. miscanthi *in China*

A total of 14520 SNPs (with less than 20% of missing data per individual and 10% per locus, Supplementary Figures S1) were obtained for the 100 individuals from the 10 Chinese populations. The levels of gene diversity (He) observed across all populations are lower than in previous studies of *S. miscanthi* in China, but similar to that of the UK population of *S. avenae* and other cereal aphids like *R. padi* (Morales-Hojas et al., 2019) (Table 2). Overall, the Chinese population is not in Hardy Weinberg Equilibrium (HWE), and the inbreeding coefficient is positive (Table 2). The same positive, significant *F*_*IS*_ is observed when we group the populations into North China (Qingdao, Tai’an, Langfang, Taigu, Pingliang and Yinchuan) and South China (Shuzou, Wuhan, Mianyang and Kunming) following the Qingling-Huaihe line (QHL; the traditional identified geographic North-South divide of China) (Table 2). Furthermore, several populations in China (Qingdao, Taigu, Yinchuan and Wuhan) of are also not in HWE, with significant, positive *F*_*IS*_ indices. Significant, positive estimates of *F*_*IS*_ is generally the result of inbreeding in the population or the effect of population subdivision, the Wahlund effect.

**Table 2.** Mean genetic diversity indices estimated for each *S. miscanthi* population, populations north and south of the QHL and each of the identified genetic clusters (GC). Ho and He are observed and expected (gene diversity) heterozygosity, respectively; *F*_*IS*_ – inbreeding coefficient.

| | He | Ho | $F_{IS}$ | $P(\text{random } F_{IS} \geq \text{observed } F_{IS})$ |
| --- | --- | --- | --- | --- |
| **Overall** | 0.28933 | 0.16720 | 0.3871 | 0.0000 |
| **North** | 0.33122 | 0.19451 | 0.37625 | 0.0000 |
| Qingdao | 0.3796 | 0.30129 | 0.16357 | 0.0117 |
| Tai'an | 0.46466 | 0.44985 | -0.01467 | 0.5147 |
| Langfang | 0.42943 | 0.40584 | 0.00621 | 0.4766 |
| Taigu | 0.2948 | 0.22396 | 0.21015 | 0.0407 |
| Pingliang | 0.41717 | 0.34735 | 0.08827 | 0.1097 |
| Yinchuan | 0.30452 | 0.22792 | 0.24868 | 0.0344 |
| **South** | 0.30482 | 0.19169 | 0.33871 | 0.0000 |
| Shuzou | 0.36834 | 0.313 | 0.10903 | 0.0689 |
| Wuhan | 0.3134 | 0.19447 | 0.36651 | 0.0004 |
| Mianyang | 0.46364 | 0.4364 | -0.00632 | 0.4827 |
| Kunming | 0.35638 | 0.33113 | 0.05435 | 0.2473 |
| GC1 | 0.46035 | 0.49644 | -0.10550 | 0.8083 |
| GC2 | 0.40616 | 0.35361 | 0.07863 | 0.0447 |
| GC3 | 0.45541 | 0.42329 | -0.01105 | 0.5731 |
| GC4 | 0.41227 | 0.37039 | 0.05263 | 0.1626 |
| GC5 | 0.36980 | 0.38917 | -0.08655 | 0.8816 |
| GC6 | 0.42755 | 0.43997 | -0.05740 | 0.5807 |

A first run of Bayesian clustering analyses of the population structure was carried out with 5 independent simulations with 100,000 burn-in and 100,000 mcmc chains for each of *K* 1–10. Analyses of the results following the Evanno method (Evanno et al., 2005) indicated that the most likely number of clusters was *K* = 6. An additional run of 5 independent simulations with 100,000 burn-in and 500,000 mcmc chains carried out for *K* 5–10 confirmed that the most likely number of *K* was 6 (Table 3, Figure 1). The structure plot shows that most sampled locations are not homogeneous, comprising individuals that are assigned to different genetic clusters (Figure 1). These results suggest that the Wahlund effect, admixture of different populations, is the most likely reason for the significant *F*_*IS*_ in Qingdao, Taigu, Yinchuan and Wuhan, which are the populations with more admixture. This is further supported by the fact that *F*_*IS*_ is non-significant when the individuals are grouped according to the genetic cluster to which they were assigned by the Bayesian clustering analyses with Structure and thus, genetic clusters are in HWE except GC2 that includes 27 individuals from seven different locations (Table 2).

**Table 3.** Table of results from Structure for the Chinese populations (5 independent simulations for K 5 – 10, 100,000 burn-in and 500,000 mcmc chains).

| K | Reps | Mean LnP(K) | Stdev LnP(K) | Ln'(K) | \|Ln''(K)\| | Delta K |
| --- | --- | --- | --- | --- | --- | --- |
| 5 | 5 | -962395 | 53822.58 | NA | NA | NA |
| 6 | 5 | -949885 | 94126.26 | 12510.06 | 49025850 | 520.852 |
| 7 | 5 | -5E+07 | 49016699 | -4.9E+07 | 55801634 | 1.138421 |
| 8 | 5 | -4.3E+07 | 59145788 | 6788294 | 28389784 | 0.479997 |
| 9 | 5 | -6.5E+07 | 41036977 | -2.2E+07 | 59738205 | 1.455717 |
| 10 | 5 | -2.7E+07 | 23812471 | 38136715 | NA | NA |

**Figure 1.**
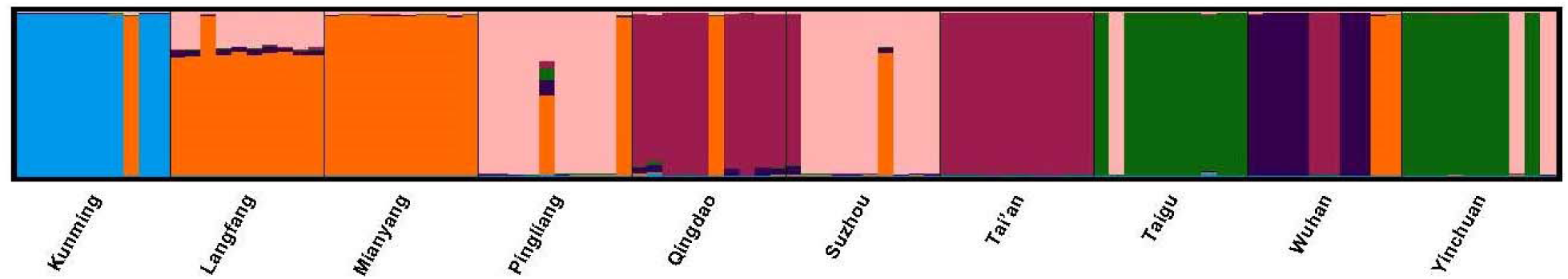
Structure analysis based on 14520 SNPs across 10 Chinese populations, with *K* = 6. Each bar represents one individual and the colours of the bars the posterior probability that each belongs to one of the six genetic clusters.

Analysis of the molecular variance (AMOVA) indicated that the overall *F*_*ST*_ of *S. miscanthi* in China was 0.3254 (*P* = 0); thus, 32.54% of the total genetic variation was explained by differences between the populations. When the individuals were grouped according to their assigned genetic cluster, 40.68% of the genetic variation (*F*_*CT*_ = 0.4069, *P* = 0) was explained by differences among the groups, indicating that the grouping of individuals from different populations but with the same genetic background increased the proportion of genetic variation explained (Table 4). Finally, we tested the QHL North-South subdivision of the *S. miscanthi* population previously suggested in the literature with an AMOVA. Results did not support the QHL divide hypothesis with only a non-significant 2.22% of the genetic variation being explained by such geographic division (*F*_*CT*_ = 0.0222, *P* = 0.2569) (Table 4).

**Table 4.**
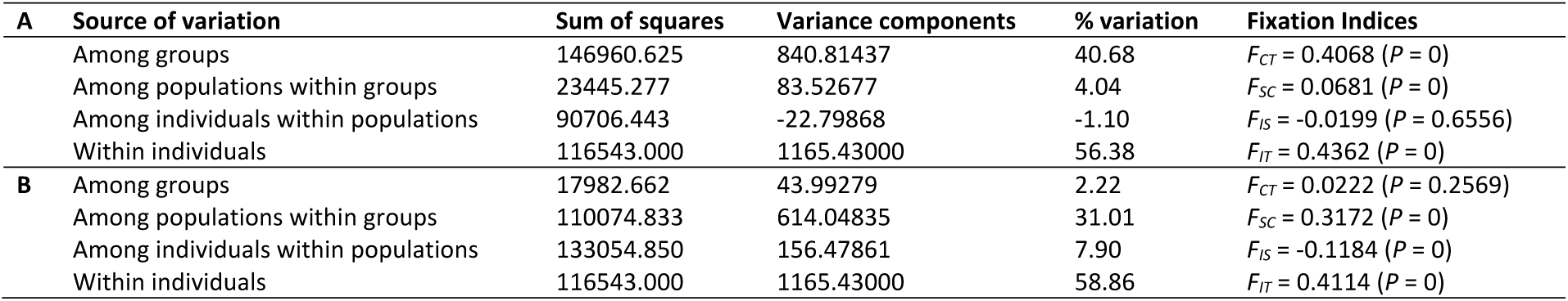
AMOVA of the SNP dataset from *S. miscanthi*. Analyses were performed to test the following hierarchical substructure: (A) individuals grouped according to the genetic cluster assignment from Structure; (B) populations grouped according to their North-South location with respect to the Qingling-Huaihe Line divide.

Pairwise *F*_*ST*_ tests showed that the genetic differentiation between the different populations is high and significant in most cases (Table 5). The only exceptions are between the populations of Suzhou and Pingliang, and Yinchuan and Taigu, which showed no genetic differentiation. Qingdao and Tai’an, which are separated by 300 kms, showed a low but significant *F*_*ST*_, and Langfang, Pingliang, Mianyang and Suzhou showed an intermediate and significant *F*_*ST*_, despite some of these locations being more than 1000 kms apart. The genetic differentiation between the genetic clusters is higher and significant in all pairwise comparisons (Table 6).

**Table 5.** Genetic differentiation between populations estimated with pairwise *F*_*ST*_ (below the diagonal) and significance (*P* values, above the diagonal) of the exact test estimated with 10100 permutations.

|  | South populations |  |  |  | North populations |  |  |  |  |  |
| --- | --- | --- | --- | --- | --- | --- | --- | --- | --- | --- |
|  | Kunming | Mianyang | Wuhan | Suzhou | Yinchuan | Pingliang | Taigu | Tai'an | Langfang | Qingdao |
| Kunming |  | 0 | 0.0001 | 0.0001 | 0 | 0 | 0 | 0 | 0.0001 | 0 |
| Mianyang | 0.38556 |  | 0.0003 | 0 | 0 | 0.0001 | 0 | 0 | 0.0001 | 0 |
| Wuhan | 0.35288 | 0.25497 |  | 0.0002 | 0 | 0.0001 | 0.0001 | 0.0003 | 0.0002 | 0.0009 |
| Suzhou | 0.35113 | 0.13993 | 0.20726 |  | 0.0004 | 0.9374 | 0 | 0 | 0 | 0.0003 |
| Yinchuan | 0.47602 | 0.36352 | 0.48536 | 0.31532 |  | 0.0012 | 0.9553 | 0 | 0 | 0 |
| Pingliang | 0.36378 | 0.12852 | 0.22950 | -0.00006 | 0.31710 |  | 0.0003 | 0 | 0 | 0 |
| Taigu | 0.52138 | 0.43957 | 0.54790 | 0.39557 | -0.01011 | 0.40377 |  | 0.0001 | 0 | 0 |
| Tai'an | 0.40604 | 0.37101 | 0.23802 | 0.29632 | 0.46615 | 0.33196 | 0.52301 |  | 0 | 0.005 |
| Langfang | 0.37918 | 0.13704 | 0.26178 | 0.11658 | 0.34895 | 0.12602 | 0.42251 | 0.35654 |  | 0.0001 |
| Qingdao | 0.34611 | 0.27895 | 0.17379 | 0.20910 | 0.40618 | 0.24716 | 0.46340 | 0.05756 | 0.27520 |  |

**Table 6.** Genetic differentiation (pairwise *F*_*ST*_) between the six genetic clusters (GC) identified with Structure. All pairwise comparisons were significant (*P* = 0) as estimated with 10100 permutations. GC 1 includes only individuals (n = 9) from Kunming; GC 2 comprises individuals from Kunming (1), Langfang (10), Mianyang (10), Pingliang (2), Qingdao (1), Suzhou (1) and Wuhan (2); GC 3 consists of individuals from Wuhan (6); GC 4 includes individuals collected in Taigu (9) and Yinchuan (8); GC 5 is formed by individuals from Qingdao (9), Suzhou (1), Tai’an (10) and Wuhan (2); and GC 6 includes individuals from Pingliang (8), Suzhou (8), Taigu (1) and Yinchuan (2).

|  | GC 1 | GC 2 | GC 3 | GC 4 | GC 5 | GC 6 |
| --- | --- | --- | --- | --- | --- | --- |
| **Genetic cluster 1** | 0.00000 |  |  |  |  |  |
| **Genetic cluster 2** | 0.42822 | 0.00000 |  |  |  |  |
| **Genetic cluster 3** | 0.46296 | 0.13319 | 0.00000 |  |  |  |
| **Genetic cluster 4** | 0.43198 | 0.30765 | 0.32792 | 0.00000 |  |  |
| **Genetic cluster 5** | 0.65910 | 0.45260 | 0.51381 | 0.54811 | 0.00000 |  |
| **Genetic cluster 6** | 0.55366 | 0.40099 | 0.44521 | 0.38811 | 0.79219 | 0.00000 |

The Mantel test was non-significant when no genetic structure within *S. miscanthi* is considered, rejecting the isolation by distance (IBD) hypothesis. However, as the Bayesian clustering analyses showed, there are 6 likely genetic clusters and this subdivision of the population can bias the test. To control for this, we performed Mantel tests for each of the identified clusters 2, 5 and 6 separately (these clusters included individuals from more than one location) but results were still non-significant. Nevertheless, the low number of individuals for some of the locations within each cluster limits the statistical power of the tests.

### *Genetic diversity and population structure of* S. avenae *in the UK*

A total of 846 SNPs with less than 25% of missing data per locus and <50% per individual (Supplementary Figure S2) were identified in 98 individuals from 12 sampling locations across England. The gene diversity, estimated as the He, is high across all populations (Table 7), although it is lower than the He observed in a previous analysis of the UK populations using microsatellites (Llewellyn et al., 2003). The Ho was higher in all populations compared to the He, resulting in high negative *F*_*IS*_ estimates which indicate that the *S. avenae* in England is not in HWE (Table 7). This contrasts with the situation of the UK population 15-20 years ago where most markers in the analysed populations were in HWE (Llewellyn et al., 2003). Although the negative *F*_*IS*_ values were not significant according to the permutation test performed by Arlequin 3.5, it should be noted that this test evaluates the probability of obtaining random values that are higher than the observed ones rather than the probability of random values being more negative. Therefore, it is most probable that the P values reported are not correct for these values. The negative *F*_*IS*_ is the result of an excess of heterozygotes, and it is considered a signature of clonal reproduction.

**Table 7.** Mean genetic diversity indices estimated for each *S. avenae* population in England and overall. N is the number of gene copies (2 x number of individuals); Ho and He are observed and expected (gene diversity) heterozygosity, respectively; *F*_*IS*_ – inbreeding coefficient. Starcross values are not included as it was represented by one single individual.

| | N | He | Ho | $F_{IS}$ | $P(\text{random } F_{IS} \geq \text{observed } F_{IS})$ |
| --- | --- | --- | --- | --- | --- |
| **Overall** | 196 | 0.28433 | 0.41243 | -0.6928 | 1 |
| Broom's Barn | 20 | 0.37289 | 0.54525 | -0.6631 | 1 |
| Hereford | 20 | 0.37214 | 0.53194 | -0.8069 | 1 |
| Kirton | 18 | 0.38078 | 0.54714 | -0.7750 | 1 |
| Newcastle | 20 | 0.38088 | 0.56614 | -0.7302 | 1 |
| Preston | 12 | 0.40722 | 0.57489 | -0.7154 | 1 |
| Rothamsted | 16 | 0.37388 | 0.51535 | -0.6035 | 1 |
| Silwood Park | 14 | 0.39285 | 0.54213 | -0.7222 | 1 |
| Starcross | 2 | - | - | - | - |
| Wye | 18 | 0.36667 | 0.51283 | -0.7137 | 1 |
| Wellesbourne | 20 | 0.35591 | 0.50472 | -0.6003 | 1 |
| Writtle | 18 | 0.36290 | 0.51392 | -0.6405 | 1 |
| York | 18 | 0.37310 | 0.53559 | -0.7419 | 1 |

Bayesian clustering analysis was run with 5 replicates of 500,000 burn-in and 1,000,000 mcmc chains to ensure convergence. Results were analysed following the Evanno method (Evanno et al., 2005), which suggested that the most likely number of genetic clusters is K = 2 (Table 8). However, this method does not estimate the deltaK for K = 1 and the Mean LnP(K) is maximised for K = 1 suggesting that the most likely number of clusters is 1 (Table 8). In addition, the standard deviation increases from K = 2, which usually happens after reaching the best K. Finally, there is no clear distribution of individuals into 2 differentiated clusters in the bar plot resulting from the analyses with k = 2, with all of them having some probability of belonging to each cluster (Figure 2b). These results suggest that there is no population structure but just one genetic cluster.

**Table 8.** Table of results from Structure for the English populations (5 independent simulations for K 1 – 12, 500,000 burn-in and 1,000,000 mcmc chains).

| K | Reps | Mean LnP(K) | Stdev LnP(K) | Ln'(K) | \|Ln''(K)\| | Delta K |
| --- | --- | --- | --- | --- | --- | --- |
| 1 | 5 | -55651.5 | 0.8927 | NA | NA | NA |
| 2 | 5 | -59437.7 | 795.9415 | -3786.2 | 4110.34 | 5.164123 |
| 3 | 5 | -67334.3 | 1512.473 | -7896.54 | 4447.6 | 2.940614 |
| 4 | 5 | -70783.2 | 4688.426 | -3448.94 | 3568.64 | 0.761159 |
| 5 | 5 | -70663.5 | 3032.39 | 119.7 | 6005.16 | 1.980339 |
| 6 | 5 | -76549 | 5716.411 | -5885.46 | 12612.16 | 2.206308 |
| 7 | 5 | -69822.3 | 2700.041 | 6726.7 | 9361.38 | 3.467125 |
| 8 | 5 | -72456.9 | 7835.535 | -2634.68 | 7552.04 | 0.963819 |
| 9 | 5 | -67539.6 | 3157.942 | 4917.36 | 4039.52 | 1.279162 |
| 10 | 5 | -66661.7 | 4041.049 | 877.84 | 2479.22 | 0.613509 |
| 11 | 5 | -63304.7 | 2208.609 | 3357.06 | 5218.48 | 2.362791 |
| 12 | 5 | -65166.1 | 3124.338 | -1861.42 | NA | NA |

**Figure 2.**
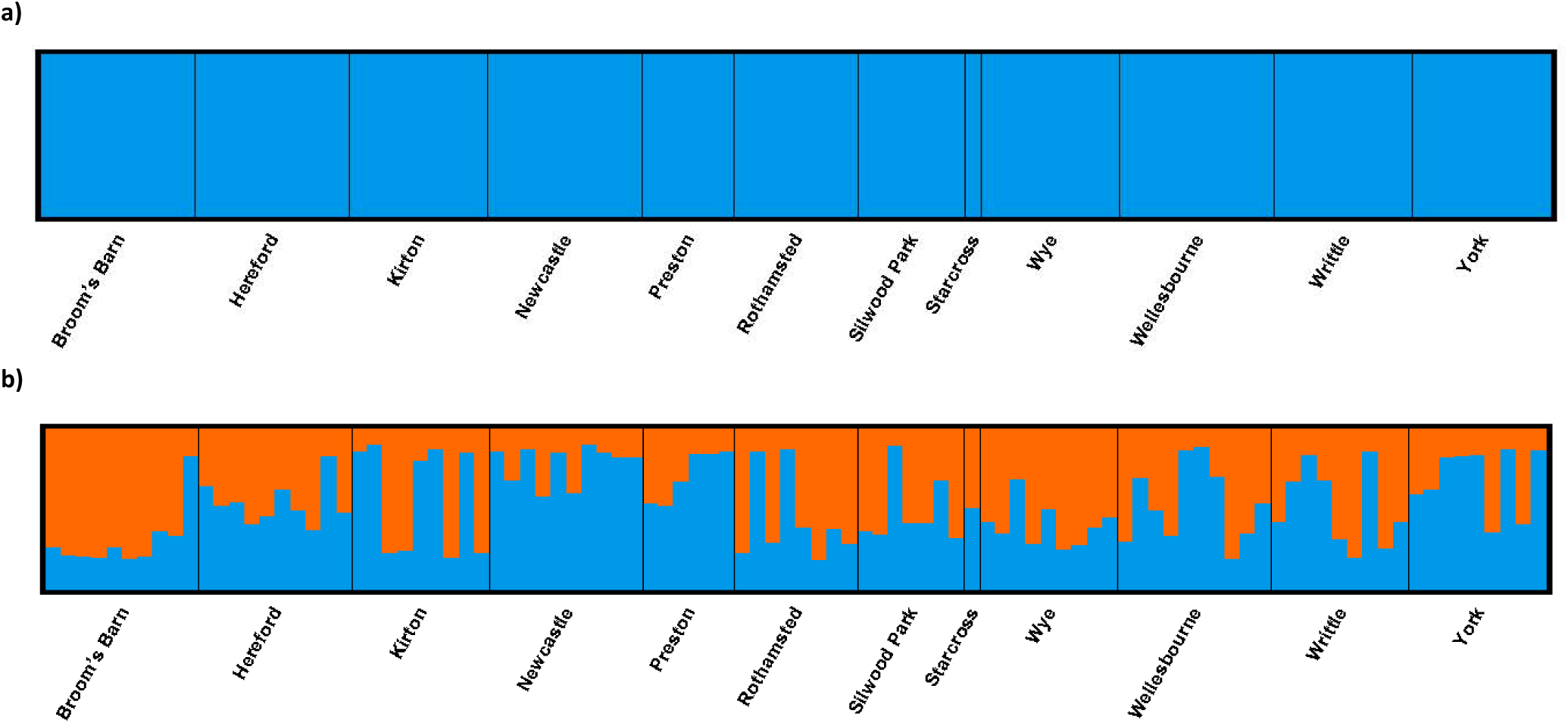
Structure analysis based on 846 SNPs across 12 English populations of *S. avenae*, with K = 1 (a) and K = 2 (b). Each bar represents one individual and the colours of the bars the posterior probability that each belongs to each of the genetic clusters.

Analyses of the Molecular Variance (AMOVA) also supported the existence of one single gene cluster. The overall diversity of *S. avenae* in England was low *F*_*ST*_ = 0.001 and non-significant among populations, indicating a low differentiation level. When the individuals were clustered according to the Structure results for K = 2, assigning individuals to each genetic cluster according to the membership coefficient estimated with clump, the amount of genetic variation that was explained between groups was 2.33% (*P* = 0) (Table 9). These results support those of Structure with one single genetic cluster and no population structure. In addition, the genetic differentiation estimates (*F*_*ST*_) were all negative (Table 10), which indicates that there is no divergence between the different locations. Similarly, when the two possible genetic clusters identified with Structure are compared, the *F*_*ST*_ is low (0.0158; *P* = 0), further supporting the lack of population structure in *S. avenae* from England.

**Table 9.** AMOVA of the SNP dataset from *S. avenae*. Analyses were performed to test the genetic cluster assignment from Structure, K = 2.

| A | Source of variation | Sum of squares | Variance components | % variation | Fixation Indices |
| --- | --- | --- | --- | --- | --- |
| | Among groups | 203.571 | 1.87014 | 2.33 | $F_{CT} = 0.0233$ ($P = 0$) |
| | Among populations within groups | 397.908 | -0.21891 | -0.27 | $F_{SC} = -0.0028$ ($P = 0.8146$) |
| | Among individuals within populations | 1765.164 | -55.77004 | -69.41 | $F_{IS} = -0.7087$ ($P = 1$) |
| | Within individuals | 13177.500 | 134.46429 | 167.36 | $F_{IT} = -0.6736$ ($P = 1$) |

**Table 10.**
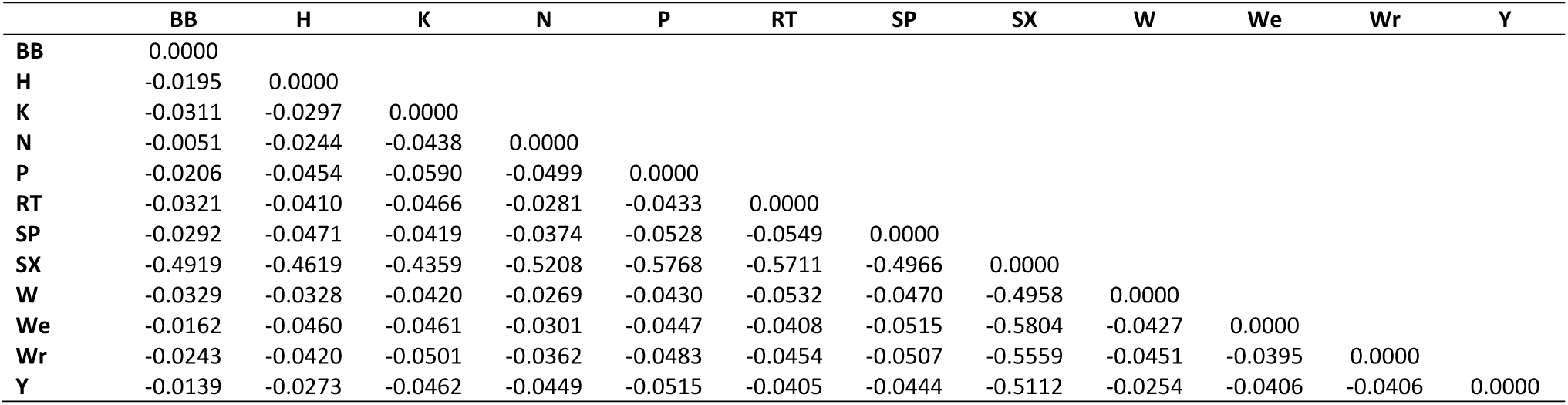
Genetic differentiation between populations estimated with pairwise *F*_*ST*_ (below the diagonal). BB – Broon’s Barn; H – Hereford; K – Kirton; N – Newcastle; P – Preston; RT – Rothamsted; SP – Silwood Park; SX – Starcross; W – Wye; We – Wellesbourne; Wr – Writtle; Y – York.

|  | BB | H | K | N | P | RT | SP | SX | W | We | Wr | Y |
| --- | --- | --- | --- | --- | --- | --- | --- | --- | --- | --- | --- | --- |
| BB | 0.0000 |  |  |  |  |  |  |  |  |  |  |  |
| H | -0.0195 | 0.0000 |  |  |  |  |  |  |  |  |  |  |
| K | -0.0311 | -0.0297 | 0.0000 |  |  |  |  |  |  |  |  |  |
| N | -0.0051 | -0.0244 | -0.0438 | 0.0000 |  |  |  |  |  |  |  |  |
| P | -0.0206 | -0.0454 | -0.0590 | -0.0499 | 0.0000 |  |  |  |  |  |  |  |
| RT | -0.0321 | -0.0410 | -0.0466 | -0.0281 | -0.0433 | 0.0000 |  |  |  |  |  |  |
| SP | -0.0292 | -0.0471 | -0.0419 | -0.0374 | -0.0528 | -0.0549 | 0.0000 |  |  |  |  |  |
| SX | -0.4919 | -0.4619 | -0.4359 | -0.5208 | -0.5768 | -0.5711 | -0.4966 | 0.0000 |  |  |  |  |
| W | -0.0329 | -0.0328 | -0.0420 | -0.0269 | -0.0430 | -0.0532 | -0.0470 | -0.4958 | 0.0000 |  |  |  |
| We | -0.0162 | -0.0460 | -0.0461 | -0.0301 | -0.0447 | -0.0408 | -0.0515 | -0.5804 | -0.0427 | 0.0000 |  |  |
| Wr | -0.0243 | -0.0420 | -0.0501 | -0.0362 | -0.0483 | -0.0454 | -0.0507 | -0.5559 | -0.0451 | -0.0395 | 0.0000 |  |
| Y | -0.0139 | -0.0273 | -0.0462 | -0.0449 | -0.0515 | -0.0405 | -0.0444 | -0.5112 | -0.0254 | -0.0406 | -0.0406 | 0.0000 |

Phylogenetic analyses were run using the two multi-SNP haplotypes of each individual. The inferred ML phylogeny comprised two major clades highly supported by bootstrap (Figure 3). Each of these two clades included one of the multi-SNP haplotypes from each individual, so that each multi-SNP haplotype is more closely related to a haplotype from another individual than to the second haplotype from the same individual. This type of phylogenetic topology can be the result of asexual reproduction from a single individual, in which all copies from each of the extant haplotypes derive from one common ancestral haplotype with no recombination. The basal node would represent the common ancestral clonal individual. Thus, the phylogeny supports the lack of population structure, and suggests that there is one single clone dominating the English population of *S. avenae*, and the asexual reproduction of this lineage.

**Figure 3.**
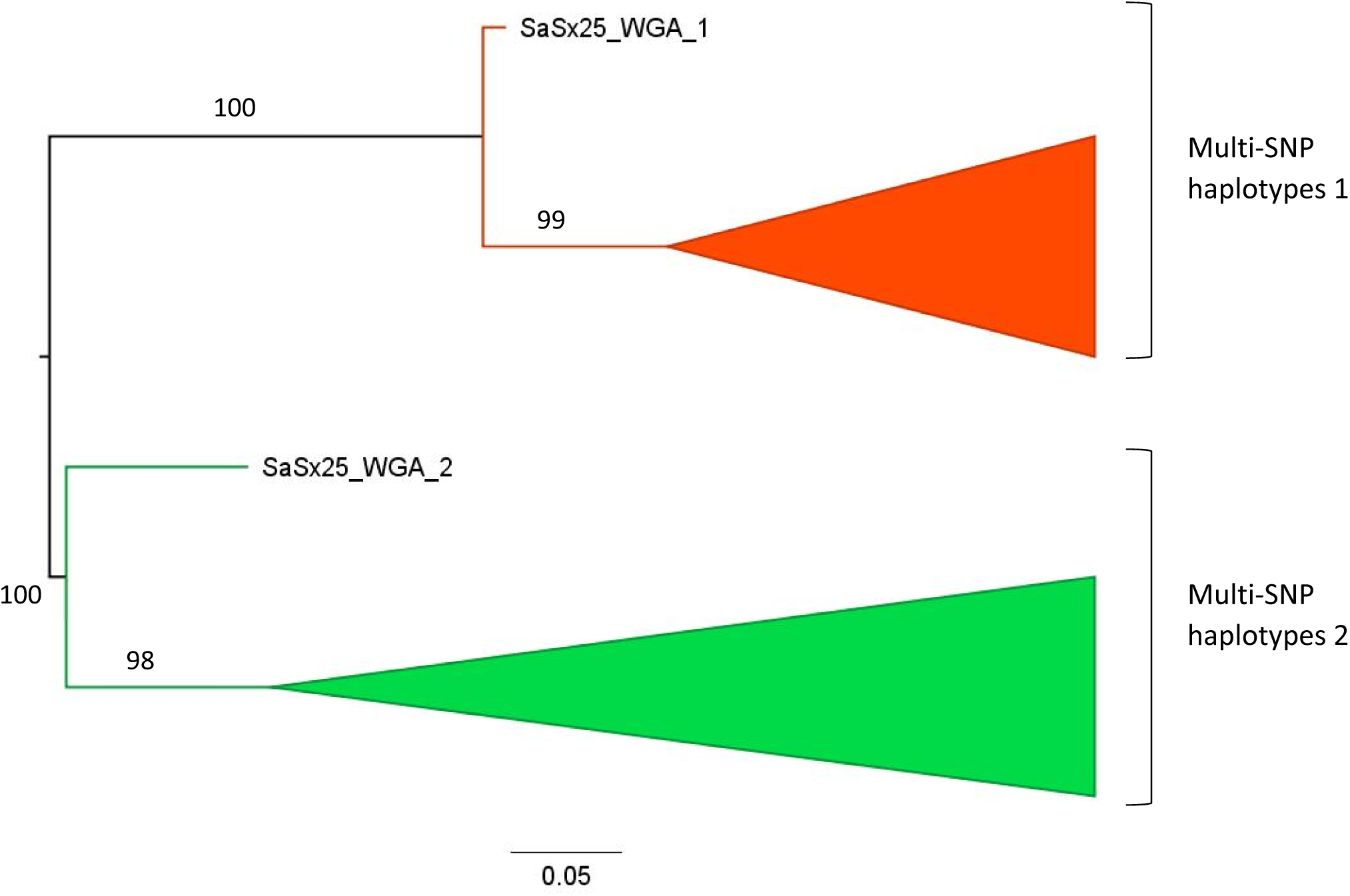
Midpoint rooted phylogenetic tree estimated with RAxML for the *S. avenae* samples from England using a dataset of 846 SNPs. The two multi-marker haplotypes from every individual are coloured in red and green, and the clades have been collapsed except for the earliest-branching haplotype of each clade. Labels on branches are bootstrap values > 90%.

Analyses of the historical demography of this species in England showed a population and spatial expansion, which would be consistent with the increase in frequency of a single insecticide-resistant clone in the population and its spread across different locations. Thus, Fu’s *F*_*s*_ index was negative (*F*_*s*_ = -25.41, *P* = 0), which is a signature of population expansions. In addition, the mismatch analyses showed a unimodal distribution and failed to reject departure from the expansion models, resulting in a non-significant Harpending’s raggedness index (0.0001, P = 1) for the demographic and spatial expansion models, and non-significant sum of squared deviation (SSD) for the spatial expansion model (P = 0.438) while significant at the 5% for the population expansion model (P = 0.031).

## Discussion

The results from the present study provides information about the evolution of two closely related species of cereal aphids under different environments and agricultural landscapes. The study demonstrates that the populations of *S. miscanthi* and *S. avenae* in China and England have evolved in the last 5 to 20 years, most likely as a result of environmental and human-induced changes such as insecticide use, and the genetic structure and diversity has changed in comparison with that observed in earlier studies. This contrasts with what it has been observed previously in another cereal aphid in England, *Rhopalosiphum padi*, whose population has not shown any change in genetic diversity or structure at least since 2003 (Morales-Hojas et al., 2019).

Results of the study indicate that *S. miscanthi* in China has a higher diversity than previously identified (Guo et al., 2005; Wang et al., 2016). This can be explained by the higher number of genome-wide molecular markers that have been used in the present analysis, but the estimated levels of genetic differentiation were similar to those of the *S. avenae* population in China (Xin, Shang, Desneux, & Gao, 2014). The species *S. avenae* is known to be present only in Yili, Xinjiang region in the northwest (Zhang, 1999), so it is likely that some previous studies of this species in China have used the incorrect taxonomic name. In addition, it could also be the case that there is undescribed taxonomic diversity within *S. miscanthi* in China, and the high levels of genetic differentiation observed could be explained by unidentified races. This is the case in Australia, where there are at least three chromosomal races (Hales, Chapman, Lardner, Cowen, & Turak, 1990), so it is possible that the situation in China is complex as well. Indeed, the high genetic differentiation observed in the present analysis between the six genetic clusters identified suggests that there are at least the same number of different taxonomic units, though the present study is not capable to determine whether they represent subspecies, host races or chromosomal races as in Australia and New Zealand.

While *S. miscanthi* in Australia and New Zealand is predominantly functional parthenogenetic (Sunnucks et al., 1996; Wilson et al., 1999), the present study indicates that the populations in China show either a heterozygote deficiency (significant, positive *F*_*IS*_) or are in HWE (not significant *F*_*IS*_), suggesting that the species reproduces predominantly by cyclical parthenogenesis. The significant deficiency of heterozygosity in populations can be explained by the admixture of different genetic clusters (Wahlund effect) observed in several populations, and the *F*_*IS*_ is not significant for the genetic clusters indicating that they are in HWE (with the exception of the GC2). This contrasts with previous studies that inferred an excess of heterozygotes in the northern populations, characteristic of anholocyclic lineages, while in the southern populations there was a deficiency of heterozygosity, which is observed in inbred sexual populations or the result of admixed populations (Wahlund effect) (Wang et al., 2016). However, it is usually the case that in colder regions populations are cyclical parthenogenetic as the aphids undergo a phase of sexual reproduction to produce eggs to overwinter; on the other hand, southern populations in warmer regimes can survive as parthenogenetic individuals throughout the year. The results of Wang et al. (2016) indicate that the contrary would be the case in Chinese *S. miscanthi*, which is unexpected. Other study of *S. miscanthi* in China showed that northern populations had heterozygote deficiency while the southern populations were in HWE or had an excess of heterozygotes, although the significance was not tested (Guo et al., 2005). The contrasting results between the present and previous analyses are also at the population subdivision. Thus, we observe no significant north-south differentiation at the QHL traditional division, identified in one previous study (Wang et al., 2016), and there is no evidence for isolation by distance (Guo et al., 2005). The population structure identified in the present study suggests that there are six genetic clusters highly differentiated. These clusters do not correspond to geographic regions and individuals from populations geographically separated by long distances can belong to the same genetic cluster; only GC1 comprising individuals from only Kunming and GC3 with individuals only from Wuhan are geographically restricted to one location. This suggests that there is long distance dispersal of *S. miscanthi* aphids across China, although the high differentiation observed between the genetic clusters suggest low interbreeding between them.

In England, the level of genetic differentiation is low across the different populations of *S. avenae* sampled and there is no evidence for genetic structure. This is in accordance to what it was previously observed (Llewellyn et al., 2003). This population homogeneity was taken to be the result of long distance dispersal of aphids, and results from the present study corroborates this. The population of *S. avenae* in England, however, has evolved in the last 15 years. While most of the markers and populations studied in 1997-1998 were in HWE, and the population showed an increase in cyclical parthenogenetic proportion with latitude (Llewellyn et al., 2003), this study shows that the present English population has an excess of heterozygotes and indicates strong clonality. This suggests that anholocycly is predominant across its range, and there is no evidence for cyclical parthenogenesis occurring towards the north of the country as expected. This change in the *S. avenae* population is most likely the result of insecticide resistance evolution. In 2011, a knockdown resistance (*kdr*) mutation to pyrethroids was detected in England’s population of *S. avenae*, and studies showed that the clone that gained this mutation spread and increased its proportion in the population from 2009 to 2014, and was also observed in Ireland from 2013 (Dewar & Foster, 2017; Foster et al., 2014; Malloch et al., 2016; Malloch et al., 2014; Walsh et al., 2019). It is therefore likely that this pyrethroid-resistant clone, which is a facultative parthenogenetic clone (Walsh et al., 2019), has continued to spread and increase in proportion, being now dominant in the English population. This is supported by the phylogenetic analysis, which shows a topology characteristic of clonal organisms. Also, the levels of genetic diversity (as measured by the He) are now lower than those of 2003 (Llewellyn et al., 2003). Although the markers used are different, it would be expected that using more, genome-wide SNPs would provide a higher level of diversity than a limited number of microsatellites. In addition, analyses of the demographic history of the population in England indicates that there has been a population demographic and spatial expansion, which substantiates the increase in proportion of the insecticide-resistant clone. Thus, as one clone gained the resistance to pyrethroids in a given location, it increased in number in the location but also expanded its distribution as it spread to other regions via migration.

Overall, this study demonstrates that the populations of two close species of cereal aphids of the genus *Sitobion* have evolved in recent years in two geographically distant regions under different environmental and human-influenced conditions. The diversity of *S. miscanthi* in China needs to be investigated more comprehensively, as the high level of genetic differentiation suggests the existence of yet unidentified forms. In contrast, the diversity of *S. avenae* has been affected by the evolution of pyrethroid resistance, and a single clone appears to be now dominating the English population. In contrast to this, *S. miscanthi* has not gained insecticide resistance despite having been subject also to its use. In England, the bird cherry – oat aphid, *R. padi*, has not evolved resistance to insecticides either, despite being sympatric with *S. avenae* and therefore subject to the same agricultural practices. Why some species evolve resistance while other do not it is still a matter of study. It has been shown in *Drosophila melanogaster* that thermotolerance influences the development and spread of insecticide resistance (Fournier-Level et al., 2019). Similarly, the distribution of cyclical and obligate parthenogenetic aphids is strongly influenced by temperature, so it is possible that there is a relationship between life-cycle type and insecticide resistance evolution in aphids. This would explain the evolution of the *kdr* and super-*kdr* mutations in the population of *S. avenae* in England, where it is predominantly anholocyclic, while the sympatric population *R. padi* and the closely related species *S. miscanthi* in China, which are predominantly cyclical parthenogenetic, have not.

## Acknowledgements

This study was made possible by a Biotechnology and Biological Sciences Research Council funded UK-China Joint Centre for Sustainable Intensification in Agriculture (CSIA) award to Rothamsted Institute BBS/OS/NW/000004, a grant from the National Key R & D Plan of China (2017YFD0201700) and the China Agricultural Science and Technology Innovation Programme (ASTIP no. CAAS-XTCX201820). We would like to thank Rothamsted Insect Survey for providing samples of *Sitobion avenae*, as part of their National Capability core activity funded by Biotechnology and Biological Sciences Research Council under the core capability grant BBS/E/C/000J0200. RMH was supported through Rothamsted’s Smart Crop Protection strategic programme BBS/OS/CP/000001, funded through the Biotechnology and Biological Sciences Research Council’s Industry Strategy Challenge Fund. We would also like to thank Thomas Mathers for providing a preliminary version of the *Sitobion avenae* genome.

## Data Accessibility

All the DNA sequencing reads have been uploaded to the European Nucleotide Archive (ENA) and can be found under the study with accession number PRJEB36151. Accession numbers for the samples’ sequencing reads are ERR3810098 – ERR3810316.

## Authors Contributions

RMH designed the project, performed the research, analysed the data, wrote and revised the manuscript.

JS collected samples, performed the research, revised the manuscript.

FAI performed the research, revised the manuscript.

XT performed the research, revised the manuscript.

JC designed the project, revised the manuscript.

