## Supplementary Tables for "Contrasting population structure and demographic history of cereal aphids in different environmental and agricultural landscapes"

**Table S1.** Description of the different filtering schemes applied to the data set. The order of rows indicates the sequential filters applied to the data. minDP – include only genotypes greater or equal to the value; minQ – include sites with quality above the value; mac – include sites with minor allele count greater or equal to the value; geno – retain sites that have been successfully genotyped in the given proportion of individuals (max-missing filtering option in vcftools); imiss – retain individuals with a proportion of missing data smaller than the value.

| FILTER | CHINESE *S. MISCANTHI* SAMPLES | ENGLISH *S. AVENAE* SAMPLES |
|---|---|---|
| **MISSING DATA** | max-missing 75 | max-missing 50 |
| **LOW-CONFIDENCE SNP CALL** | minDP 3<br>mac 3<br>minQ 30 | mac 3<br>minQ 30<br>minDP 3 |
| **MISSING DATA** | remove-indels | max-missing 50<br>imiss > 50%<br>max-missing 75<br>remove-indels |
| **INFO FILTERS** | thin 2000 | thin 2000 |
| **MISSING DATA** | max-missing 0.90 | |
| **INFO FILTERS** | thin 5000 | |
| **ALL SAMPLES SNPS** | 14520 | 846 |
| **ALL SAMPLES INDIVIDUALS** | 100 | 98 |

Initial dataset of 100 individuals and 564295 SNPs for Chinese *S. miscanthi*.
Initial dataset of 119 individuals; 2248285 SNPs for English *S. avenae*.

**Table S2.** Geographic distances between the different locations sampled in Chin. The distances are in kilometres (Kms) and have been estimated in straight lines in Google maps.

| | Yinchuan | Langfang | Pingliang | Qingdao | Tai'an | Taigu | Kunming | Mianyang | Suzhou | Wuhan |
|---|---|---|---|---|---|---|---|---|---|---|
| **Yinchuan** | 0 | | | | | | | | | |
| **Langfang** | 934 | 0 | | | | | | | | |
| **Pingliang** | 348 | 980 | 0 | | | | | | | |
| **Qingdao** | 1279 | 508 | 1242 | 0 | | | | | | |
| **Tai'an** | 990 | 390 | 947 | 300 | 0 | | | | | |
| **Taigu** | 564 | 450 | 550 | 704 | 426 | 0 | | | | |
| **Kunming** | 1546 | 2120 | 1287 | 2095 | 1875 | 1668 | 0 | | | |
| **Mianyang** | 792 | 1430 | 534 | 1540 | 1276 | 980 | 750 | 0 | | |
| **Suzhou** | 1537 | 980 | 1394 | 530 | 666 | 1010 | 1872 | 1507 | 0 | |
| **Wuhan** | 1156 | 1015 | 940 | 827 | 710 | 785 | 1287 | 925 | 598 | 0 |
