## Supplementary Figures for "Contrasting population structure and demographic history of cereal aphids in different environmental and agricultural landscapes"

**Figure S1.** Proportion of missing data per individual (a) and per locus (b) in the Chinese *S. miscanthi* SNP dataset after applying the filtering scheme.

a)

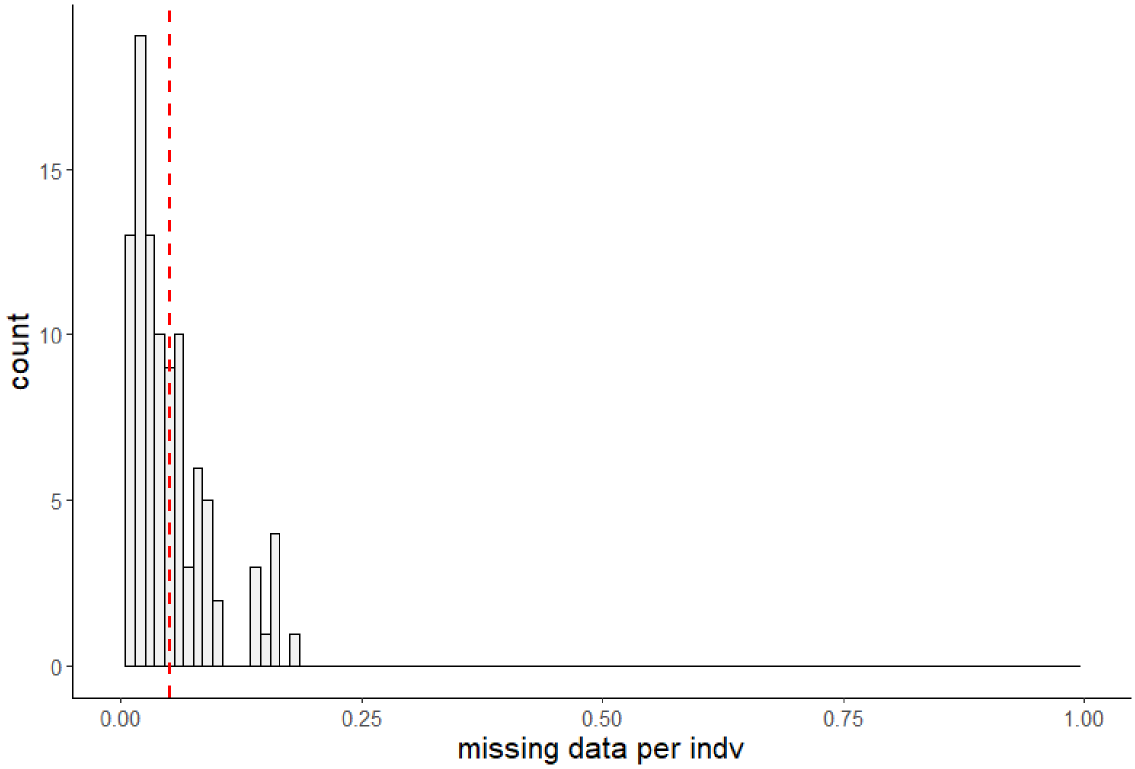


b)

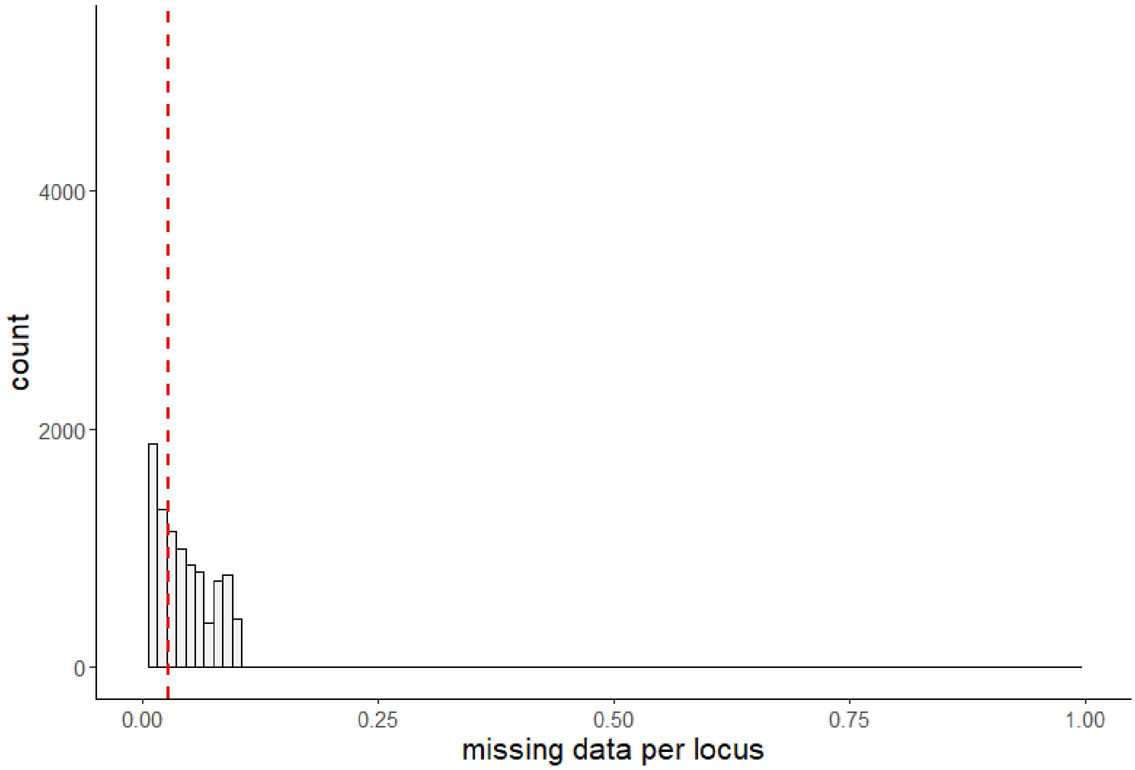

**Figure S2.** Proportion of missing data per individual (a) and per locus (b) in the English *S. avenae* SNP dataset after applying the filtering scheme.

a)

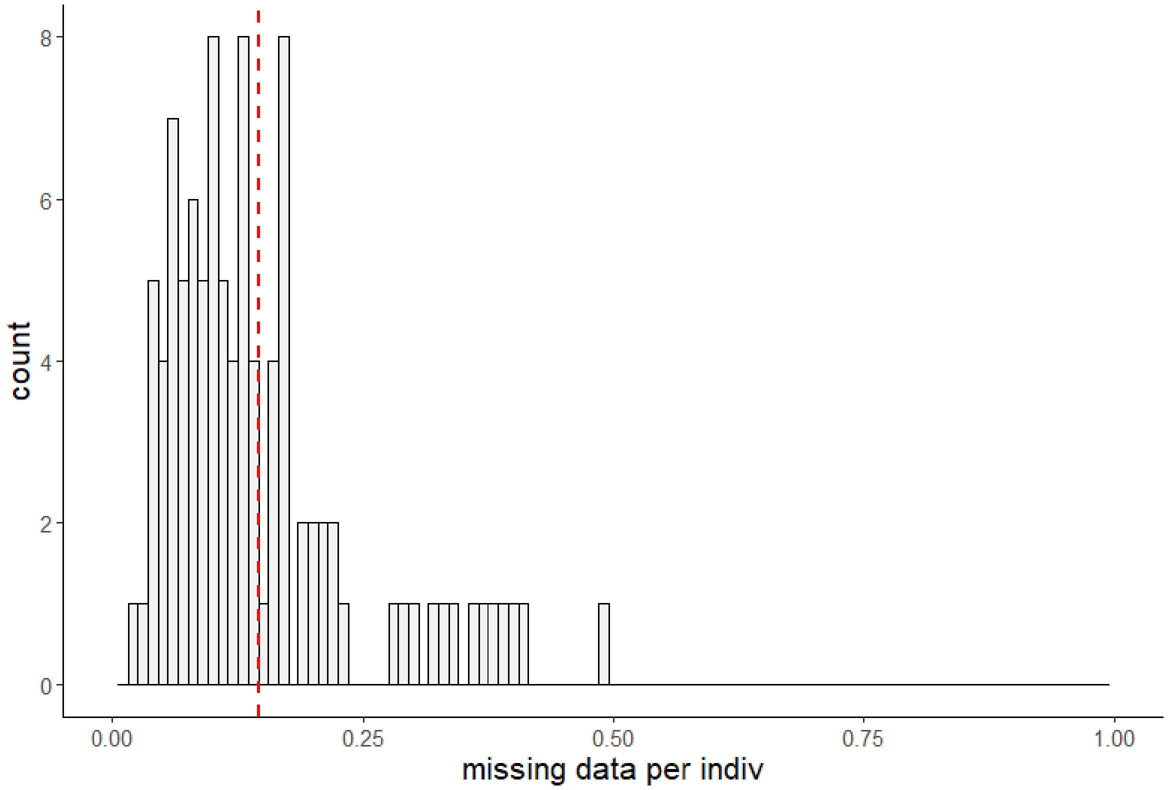


b)

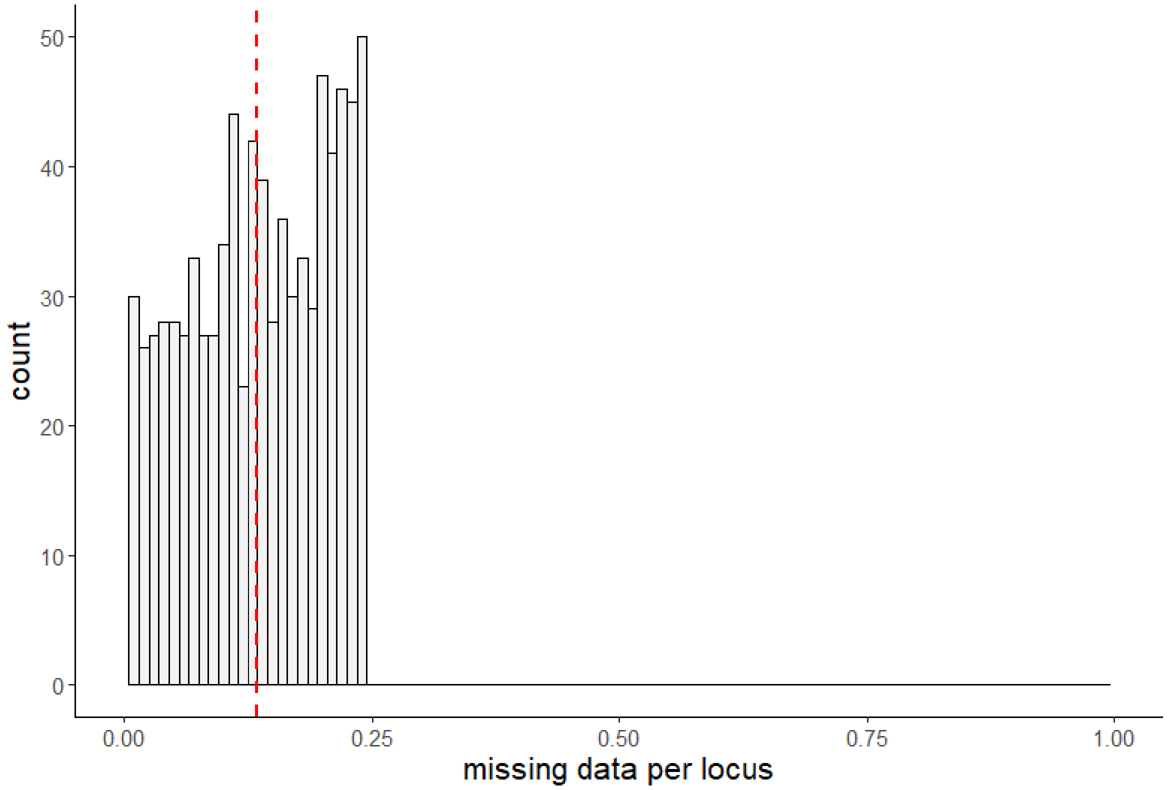
